# Rapid discovery of new-to-nature protein domains by novelty-first forcing of language models

**DOI:** 10.1101/2025.10.02.679910

**Authors:** Arjuna Subramanian, Matt Thomson

**Affiliations:** Division of Biology and Biological Engineering, California Institute of Technology, Pasadena, CA 91125

## Abstract

Approximations for the existence and extent of physically permissible protein structures beyond those found in nature vary wildly. As predicted structure databases swell thanks to abundant sequence data and generative protein design models concurrently grow in their power to propose new aspects of protein structure, these questions and those of which essential features (e.g. stability, function, robustness) distinguish natural domains from novel ones have been cast in even sharper relief. We demonstrate that protein language models (PLMs) can simultaneously innovate in sequence and structure to suggest new-to-nature protein domains displaying supersecondary and tertiary elements outside of categorized CATH superfamilies. Developing and applying two orthogonal processes for obtaining compact and globular folds from PLMs without bias from other physicochemical or functional constraints, we discover putative novel domains that emerge parallel to known natural ones at rates far exceeding those obtainable by bioinformatic mining of structure databases. Computational characterization of these domain candidates indicates that many exhibit reasonable folding thermodynamics and kinetics, suggesting that natural protein structure-space is far from biophysically complete. These results point away from stability as the definitive selective force behind the observed landscape of real protein folds, and insinuate that many unrealized folds may be equally consistent with the structural rules of protein-based life.

## 1 Introduction

At the center of the pursuit of novel protein structures lies a fundamental biophysical mystery – why might some (or many) compact three-dimensional structures be permitted by the laws of physics and yet be apparently unrealized in nature? Perhaps these hypothetical domains are encoded by a relative dearth of fold-encoding sequences, want for robustness to mutation, and were lost to drift [Govindarajan and Goldstein, 1996, Helling et al., 2001, Li et al., 1996, Bornberg-Bauer, 1997]. Or they lack functional fitness, particularly the metal-scaffolding and RNA/DNA-binding modalities thought to predominate among the first proteins, and were casualties of selection [Dupont et al., 2010, Vyas et al., 2021]. Maybe, on the other hand, flaws in folding thermodynamics or kinetics disfavor or even outright forbid them, and the documented set of structural units is complete after all [Watters et al., 2007, Baker, 2000, Yue and Dill, 1995].

Opinions on the structural completeness question have historically been sharply split [Taylor et al., 2009, Chitturi et al., 2016, Zhang et al., 2006, Skolnick et al., 2012]. Physics-based models have yielded a small handful of *de novo* designed proteins with truly new-to-nature folds [Kuhlman et al., 2003, Minami et al., 2023, Sakuma et al., 2024]. Converting protein structure prediction models to directed hallucination and diffusion models has accelerated the unconditional generation of backbones, although it remains unclear whether these are better described as novel folds or as stretching the fringes of extant folds [Anishchenko et al., 2021, Watson et al., 2023]. The advent of massive-scale predicted structure databases, dwarfing the ∼200, 000 experimental structures of the Protein Data Bank (PDB) with a rich trove of nearly 1 billion AlphaFold, ESMFold, and ColabFold predictions has further muddied views on the completeness of natural structures [Varadi et al., 2022, Lin et al., 2023, Kim et al., 2025a,b]. The structural “dark clusters” and newly collated Pfam families and CATH superfamilies carved from these databases make the PDB, at least, look incomplete indeed [Barrio-Hernandez et al., 2023, Pavlopoulos et al., 2023, Durairaj et al., 2023, Lau et al., 2024]. In quantitative terms, a ∼ 1000x inflation of individual structures (the UniRef50 portion of the AlphaFoldDB) was shown to boost the number of CATH superfamilies from 5,841 to 6,573 (level “H”, +12.5%), and the number of CATH topologies/folds from 1,349 to 2,081 (level “T”; +54.3%), good for a new topology discovery rate of 732*/*214, 683, 839 ≈ 3.4 × 10^6^, or 3-4 per every one million newly predicted structures [Lau et al., 2024]. Despite this expanded accounting and classification of folds, the original question persists – given the collection of natural protein domains, experimentally-determined *and* model-predicted combined – how much can nature really have left by the wayside as far as stable protein topologies? And as a practical matter, how can these never-realized structural motifs be readily discovered without sifting through millions of new predictions or pre-specifying possible atomic coordinates and properties *a priori*?

On the surface, protein language models (PLMs) may seem an odd choice of tool for unearthing novel structures, given that they are explicitly sequence models. It is well-established, however, that that transformer-based PLMs implicitly capture key features of structure, function, and even evolutionary history in the process of learning sequence features and constraints [Chen et al., 2023, Hayes et al., 2024, Hie et al., 2022]. Generative PLMs can make small structural and/or functional changes, for example, by infilling nature-adjacent variants into an enzyme class [Munsamy et al., 2022, Madani et al., 2023]. They can also access putatively novel structures without natural homologs albeit at low frequencies or after significant optimization – through multiple free generation schemes [Ferruz et al., 2022, Verkuil et al., 2022]. We reason that unlocking the full latent capacity of PLMs to access new domain structures requires targeted strategies to enrich for rare pinpricks of structural novelty in generative output, without introducing functional biases. Thus, we build and deploy two distinct fitness-agnostic strategies – one inspired by fold recombination events in real-world protein evolution, the other a direct selection against resemblance to natural motifs – that drive PLMs towards novel structure generation. These complementary approaches deliver an abundance of novel domains computationally projected to be stable, foldable, and un-mappable to any CATH example, illustrating multiple sequence-centric routes to expanding protein structural diversity and suggesting that the known structural catalog is far from complete with respect to the governing biophysics.

## 2 Results

### 2.1 Novel domains emerge from a fold-recombining genetic algorithm

One potential avenue for finding novel protein domains is to start from primitive structural elements and recombine them, evolve them, and put them under selective pressure, all in *in silico*. This approach is a genetic algorithm for domain diversification, loosely inspired by hypotheses for how early enzymes and ancient protein folds may have originated from fused and mutated primordial polypeptides exploring new folding landscapes. [Longo et al., 2020, Alvarez-Carreño et al., 2022]. As starting material to seed the algorithm, we generate a small library of 800 miniprotein-sized (length: 40 amino-acids) fragments *de novo* via PLM-informed replica-exchange Markov Chain Monte Carlo sampling. Briefly, random amino-acid sequences are evolved in single mutation steps subject to an energy function that favors greater sequence likelihood and structural contact density, both as inferred by ESM2-650M (refer to full implementation details in Section 4.1.1). The resulting miniproteins sample variable *α* and *β* secondary-structure content, loop size, packing geometry, super-secondary organization, and degrees of (dis)order (Fig. S.1). The choice of *de novo* generation is motivated by a desire to mitigate against sequence-side biases in favor of nature that might be introduced by alternative such as fragmenting real or experimental structures from published databases. Indeed, while structure-based search with Foldseek (504*/*800 = 63.0% hit rate against the UniRef50 portion of the AlphaFoldDB) shows that the generated fragments are plausible and representative building blocks, sequence-based search with MMseqs2 (48*/*800 = 6.0% hit rate against UniRef50) indicates that they are distinct from natural sequences, both as desired.

We carry forward a randomly selected subset of 100 mini-protein fragments as the initial population for the genetic algorithm, which proceeds for 200 epochs. In each epoch, 20 recombined and mutated fragments are generated and evolved over the same energy landscape as used for the fragment library before being added to the population; stochastic selection with survival rate proportional to the fraction of total linear-chain amino-acid surface-area buried (“burial fraction”) is applied to reduce the population back to a constant target size of 100 (refer to Section 4.1.2 for full implementation details, including the form of the selection function). Burial fraction is a function-unaware selective force that rewards compactness and globularity, allowing stable tertiary folds, alike-to-nature and new-to-nature both, to emerge. The mean burial fraction increases with time, demonstrating that compact folds become more common and/or folds become more compact on average as the algorithm proceeds (Figure 1A). Assigning CATH labels wherever possible with a Foldseek-based annotation pipeline, natural folds accrue at a roughly constant rate of 2.4 per epoch, while compact (thresholded at burial fraction *>* 0.5) yet novel folds emerge sporadically; the first new-to-nature fold (ID: 3733B8_R10) appears in epoch 10 with subsequent interfold arrival times as long as 45 and as short as 2 epochs (Figure 1A-B). Working off of building blocks that are almost exclusively displaced from nature in sequence but nearby in structure, the algorithm reaches ∼500 natural folds and 15 putatively novel ones, suggesting that addenda to natural structure-space are surprisingly accessible to a novelty-prioritizing search that does not specify backbones *a priori*.

**Figure 1.**
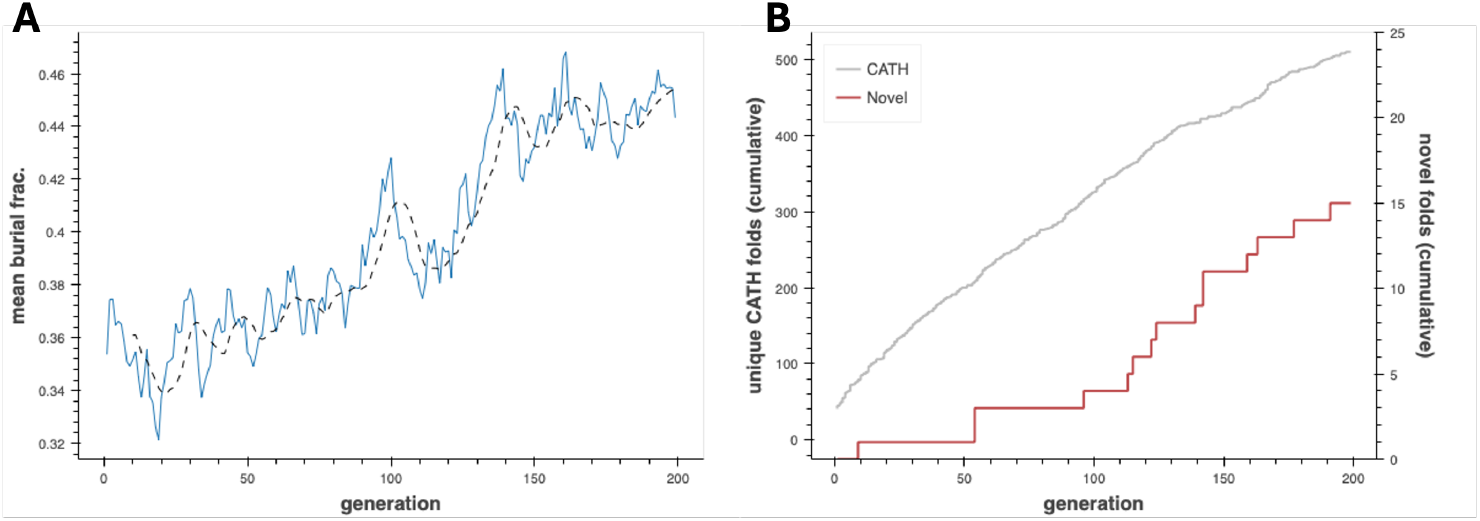
Emergence of novel folds from a PLM-based genetic algorithm. (**A**) Mean fractional amino-acid surface burial (protein compactness proxy) over 200 generations of the structure discovery genetic algorithm. (**B**) Cumulative counts of unique CATH-annotated folds and putative novel folds detected over 200 generations of the structure discovery genetic algorithm.

The 15 new-to-nature domains proposed by the evolutionary algorithm are markedly distinct from their nearest CATH analogs and structurally diverse, visiting three of the four major topology classes – all-*α* (120FD5_R127, 244D7D_R143, 794026_R125, A49A4F_R116, A783532_R160, B4FC4F_R164, BC2987_R55, DB6817_R178, F99539_R114), all-*β* (0CF85E_R97, 3733B8_R10, C86FA9_R143), and *α* + *β* (120FD5_R127, 9D1265_R55, A0A7B8_R123) (Fig. 2, Table S.1). It is curious that no novel *α/β* folds are observed, given the prominent functional speciation of such domains in nature [Choi and Kim, 2006].

**Figure 2.**
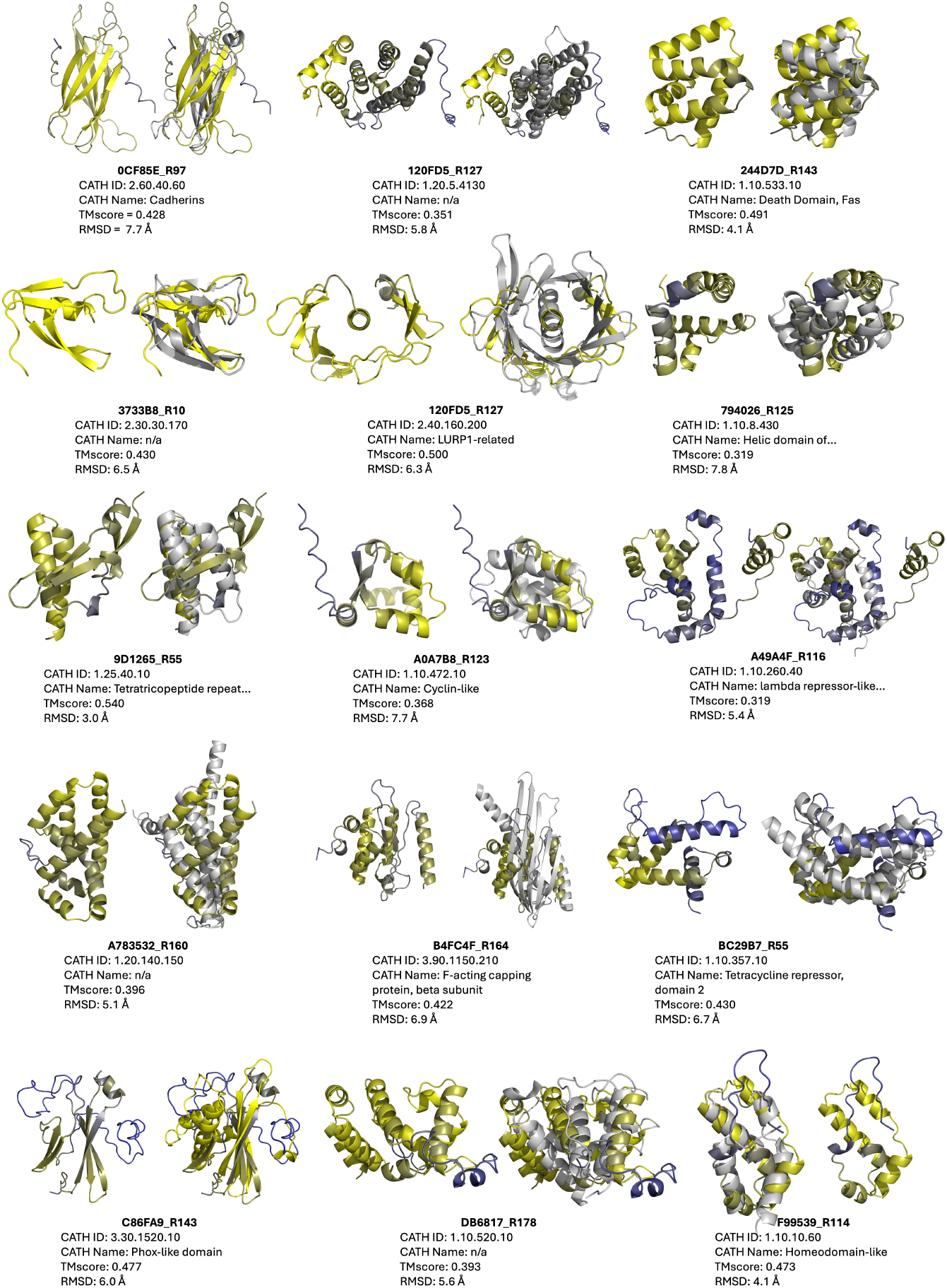
Fifteen novel folds achieved by the structure discovery genetic algorithm. Within each pair: **left** – putative novel fold (colored by ESMFold pLDDT; yellow=high, blue=low); **right** superimposed with closest CATHDB50 Foldseek hit in TMalign mode, with CATH metadata and global alignment metrics reported below.

### 2.2 “Inverse-folding funnels” distinguish putatively stable folds

For additional insight into this handful of novel domains and whether they are truly plausible as far as the thermodynamics and kinetics of protein folding, we introduce the “inverse-folding funnel” as a heuristic for computationally assessing the stability of a putative novel fold. This exercise is roughly analogous to the use of Rosetta *ab initio* structure prediction simulations to explore a protein-folding energy landscape [Minami et al., 2023]. Traditionally, if a plot of estimated energy vs. backbone RMSD vs target for many replicates of the same sequence, indicates: (1) an association between lower energy (higher stability, i.e. favorable folding thermodynamics) and smaller RMSD; and (2) an absence of “trapped” subpopulations at moderate-to-high RMSD and local energy minima (presumed metastable states, indicators of poor folding kinetics), then it resembles the prototypical folding funnel of a globular protein spontaneously collapsing to its native-state structure, whereas one failing either or both criteria warns of folding pathologies precluding viable expression [Dill and Chan, 1997]. We instead use an inverse-folding model (ProteinMPNN from Dauparas et al. [2022]) to generate many sequence-diversified versions expected to encode each of the novel domain structures from Figure 2, provided as backbone templates. Preclustering by sequence similarity to minimize redundancy, we predict structures with ESMFold, estimate absolute energies with Rosetta, and quantify global alignment between inverse-folded structures and templates as TMscores. For the “inverse-folding” version of the funnel, we look for correlation between lower energy and *higher* TMscore and for a lack of low-TMscore/low-energy states – the former remains a proxy for thermodynamic stability, while the latter rules out metastable intermediates and the distant possibility that a “novel” domain might be an noised version of a CATH domain recoverable by the slight re-noising applied in inverse-folding.

For 8 of the 15 putatively novel domains,(IDs: 0CF85E_R97, 244D7D_R143, 26D32B_R192, 3733B8_R10, A49A4F_R116, A78532_R160, B4FC4F_R164, DB6817_R178), this procedure evinces a convincing funnel with the aforementioned essential characteristics, bolstering confidence that these are realizable new-to-nature structures (Fig. 3). Other faux folding landscapes point to problem spots; for example for 9D1265_R55 multiple equivalent energy minima are observed, while for BC29B7_R55 a single minimum is centered around a TMscore well less than 0.5, as if inverse-folding reliably converges to a more stable neighbor in structure-space (Fig. 3). Other landscapes are more flat than funnel-shaped, as in the case of F99539_R114, implying that some novel domain candidates may lack a true native state. As general guidelines for stable and robust structures, we additionally set rough threshold values of *<* −2.2 REU/aa and TMscore *>* 0.5 for inverse-folded variants to clear and note that even for those domains that do exhibit funnel-like folding landscapes many variants can fail one or both, reiterating the importance of re-noising for recovering more-plausible adjacent structures from novel domain candidates. Although not all potentially novel folds are created equal as far as presumed folding dynamics and stability, fold recombination and evolution from artificial fragments inculcates a strong belief that novel domains beyond natural structure-space are readily accessible in the absence of function-centric selection.

**Figure 3.**
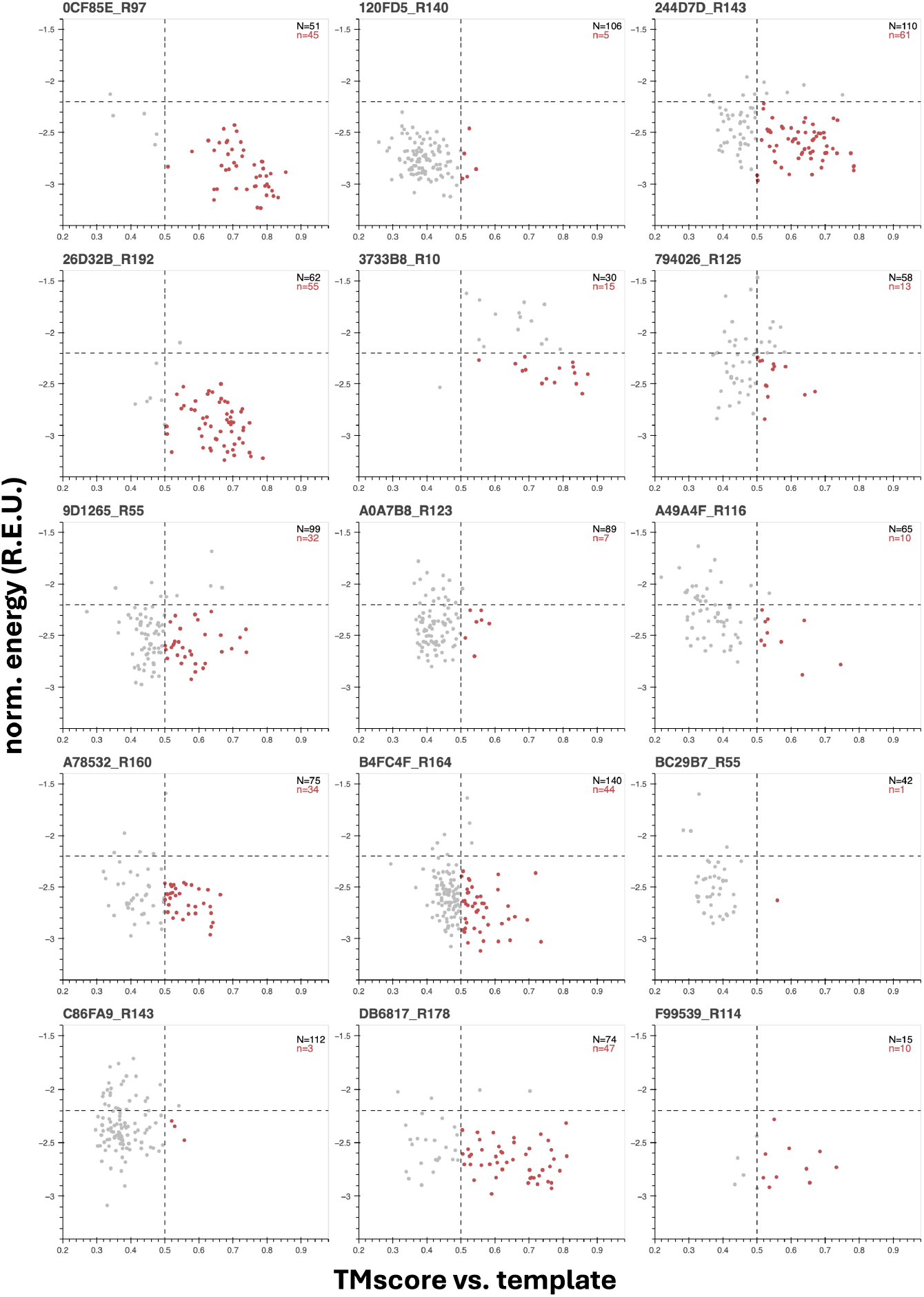
“Inverse-folding landscapes” for fifteen novel folds achieved by the structure discovery genetic algorithm suggest variable stability. Length-normalized energies (from Rosetta) vs. TM-score (from Foldseek in TMalign mode) for ProteinMPNN-designed sequences inverse-folded off of structure discovery genetic algorithm putative novel folds as templates. Gray dots correspond to all sequences/structures for a given template after clustering 200 initial sequences per template at 60% sequence similarity. Red dots show the subset of inverse-folded seqUences whose ESMFold-predicted structures pass an energy scoring threshold (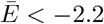 REU/aa) and the standard TM-score global match threshold (TMscore *>* 0.5).

### 2.3 Structure-first foldtuning enriches for domains with new-to-nature structures

Although the fold recombination genetic algorithm recovers novel structures, these new-to-nature folds emerge at a ∼30x slower rate compared to natural ones. We reasoned that a more rapid and direct route to novel domains might be to transform PLM foldtuning – a sequence-perturbing, fold-preserving method for novel sequence discovery – into a fold-perturbing, sequence-insensitive method for novel structure discovery [Subramanian et al., 2025]. We first estimated the latent capacity of ProtGPT2, the default PLM of foldtuning, to generate previously unseen structural motifs off-the-shelf without additional training. To do so, we revisited a dataset of ∼3 million ESMFold-predicted small protein structures obtained by autoregressive free sequence generation from ProtGPT2 across thirty (top_k, temperature) sampling hyperparameter pairs. We downsampled this dataset by 10x stratifying by hyperparameter pair, annotating with CATH labels wherever possible via our Foldseek-based pipeline. Compactness/globularity was estimated and reported via the burial fraction, as before. Aggregated results are reported in Table 1. As thresholds for putative novel structures, we look for predicted structures with a fractional burial *>* 0.5 and no assignable CATH domain label; occurrence rates range from 0.11% for top_k 1500 and temperature 0.8 to 0.41% for top_k 4000 and temperature 5.0. In general, increasing either hyperparameter corresponds to an increase in this novelty rate, but this trend is not monotonic. Depending on whether structural diversity is defined at the CATH superfamily level or CAT fold/topology level, structural diversity expands or contracts, respectively, with increasing top_k and/or temperature. This implies that hyperparameter selection associated with sequence novelty (“textual novelty”) favors finer-grained structure perturbations typical of superfamily-to-superfamily variation over supersecondary rearrangements indicative of satisfyingly novel *folds*. The fraction of compact proteins generated (burial fraction *>* 0.5) also consistently drops by roughly 2x as temperature goes from 0.8 to 5.0. Accordingly, to strike a balance between compactness, CATH non-assignability, and the magnitude of structural perturbation, we fix sampling hyperparameters at top_k 950 and temperature 1.5 for what we refer to as “structure-first” foldtuning.

**Table 1:** CATH domain coverage, structural compactness, and novel fold discovery rate from base ProtGP2 sampling hyperparameter scan. CAT(H) folds(superfamilies) detected, CATH hit absence (no hit with TMscore *>* 0.5), structural compactness (burial fraction *>* 0.5), and novel fold discovery rate for 30 sampling hyperparameter combinations from varying top_k (vocabulary size: 600, 950, 1500, 2400, 5000) x temperature (0.8, 1.0, 1.2, 1.5, 2.0, 5.0).

| Hyperparams |  | Results |  |  |  |  |
| --- | --- | --- | --- | --- | --- | --- |
| top_k | temp | # CATH | # CAT | No CATH | Compact | Both |
| 600 | 0.8 | 905 | 401 | 0.224 | 0.2376 | 0.0016 |
|  | 1.0 | 935 | 412 | 0.2121 | 0.2298 | 0.0021 |
|  | 1.2 | 975 | 404 | 0.2189 | 0.2176 | 0.0025 |
|  | 1.5 | 988 | 408 | 0.2218 | 0.1983 | 0.0023 |
|  | 2.0 | 977 | 407 | 0.2418 | 0.1839 | 0.0025 |
|  | 5.0 | 967 | 384 | 0.3628 | 0.1039 | 0.0017 |
| 950 | 0.8 | 908 | 402 | 0.2143 | 0.2361 | 0.0018 |
|  | 1.0 | 955 | 416 | 0.2115 | 0.2293 | 0.0018 |
|  | 1.2 | 988 | 430 | 0.2261 | 0.1996 | 0.0031 |
|  | 1.5 | 984 | **419** | 0.2347 | **0.1922** | **0.0037** |
|  | 2.0 | 994 | 421 | 0.2432 | 0.1746 | 0.0036 |
|  | 5.0 | 996 | 394 | 0.3584 | 0.1008 | 0.0029 |
| 1500 | 0.8 | 954 | 404 | 0.2145 | 0.2313 | 0.0011 |
|  | 1.0 | 964 | 410 | 0.2228 | 0.2113 | 0.0029 |
|  | 1.2 | 994 | 418 | 0.2378 | 0.1908 | 0.0023 |
|  | 1.5 | 1014 | 415 | 0.2464 | 0.1727 | 0.0028 |
|  | 2.0 | 1005 | 403 | 0.2612 | 0.1528 | 0.0028 |
|  | 5.0 | 1017 | 382 | 0.3634 | 0.095 | 0.0028 |
| 2400 | 0.8 | 941 | 406 | 0.2227 | 0.2221 | 0.002 |
|  | 1.0 | 970 | 410 | 0.2279 | 0.2045 | 0.002 |
|  | 1.2 | 993 | 420 | 0.247 | 0.1804 | 0.0029 |
|  | 1.5 | 1025 | 412 | 0.2572 | 0.1582 | 0.0036 |
|  | 2.0 | 1055 | 425 | 0.2734 | 0.1417 | 0.0034 |
|  | 5.0 | 1054 | 396 | 0.3536 | 0.0963 | 0.0033 |
| 4000 | 0.8 | 962 | 433 | 0.2232 | 0.2303 | 0.0024 |
|  | 1.0 | 1021 | 440 | 0.2183 | 0.2001 | 0.0026 |
|  | 1.2 | 1012 | 418 | 0.2521 | 0.1767 | 0.0022 |
|  | 1.5 | 1076 | 425 | 0.2539 | 0.1519 | 0.0023 |
|  | 2.0 | 1010 | 380 | 0.2786 | 0.1358 | 0.0027 |
|  | 5.0 | 1008 | 390 | 0.341 | 0.1028 | 0.0041 |

Structure-first foldtuning (refer to 4.1.3 for full details) proceeds for five rounds. In a given round, 10,000 sequences are generated out of the current (*k*-th) model and filtered based on predicted structures to enforce compactness (burial fraction *>* 0.5) and CATH non-assignability (no Foldseek-TMalign hit in CATHDB50 with TMscore *>* 0.5). Filtered sequence-structure pairs are ranked in order of descending burial fraction, with the 100 most-compact becoming the training set used to finetune the (*k* + 1)-th model. Over five rounds, structure-first foldtuning progressively enriches for sequence-structure pairs meeting the compactness/non-assignable novelty criteria, from 111/10,000 (11.1%) after one round to 269/10,000 (26.9%) after five (Table 2). Neither burial fraction nor the number of unique CATH domains is observed to change significantly at the population level, with a concomitant drop in the CATH assignability rate across all sequences/structures, a further indication that while a non-globular sub-population persists, all of the growth in structural diversity is diverted to putatively novel domains.

**Table 2:** Emergence of novel and CATH-annotated domains over five rounds of “structure-first” foldtuning. Number of generated sequences successfully annotated with a CATH domain by Foldseek (“# CATH”), structural hit rate (fraction of generated sequences assigned to *any* CATH label), number of generated sequences assigned as putative novel folds (“# Novel”; burial fraction *>* 0.5 and no hit with TMscore *>* 0.5), and mean burial fraction over the course of five rounds of structure-first foldtuning with top_k 950, temperature 1.5, and 10,000 sequences sampled per round.

| Round | Mean Burial Frac. | # Novel | # CATH | Struct. Hit Rate |
| --- | --- | --- | --- | --- |
| 1 | 0.433 | 111 | 1166 | 0.719 |
| 2 | 0.444 | 192 | 1190 | 0.641 |
| 3 | 0.438 | 206 | 1178 | 0.589 |
| 4 | 0.451 | 240 | 1155 | 0.632 |
| 5 | 0.438 | 269 | 1171 | 0.549 |

Structure-first foldtuning proposes 1018 novel domains in total over five rounds, a remarkable rate of 2.0% of all generated proteins. To check for redundancy, this set of 1018 is clustered at a TMscore*>* 0.5 global alignment threshold, consistent with grouping templates that would occupy the same superfamily and/or fold if added to the CATH database; this marginally reduces the number of domains to 916. Even applying a stricter structural novelty criterion – ruling out any example with TMscore*>* 0.5 to any domain in the entire AlphaFoldDB50 database, not merely its CATHDB50 subcomponent – only contracts this set to 762 members in total. We identify a final high-priority set of putative novel domains by ranking in order of descending burial fraction and taking the top 100 most-compact. Inverse-folding landscapes are constructed as for the genetic algorithm outputs. Predicted structures (with and without closest CATH hits) and inverse-folding landscapes for the best 10 templates as ranked by average estimated folded-state energy are shown in Fig. 4 and Fig. 5 respectively.

**Figure 4.**
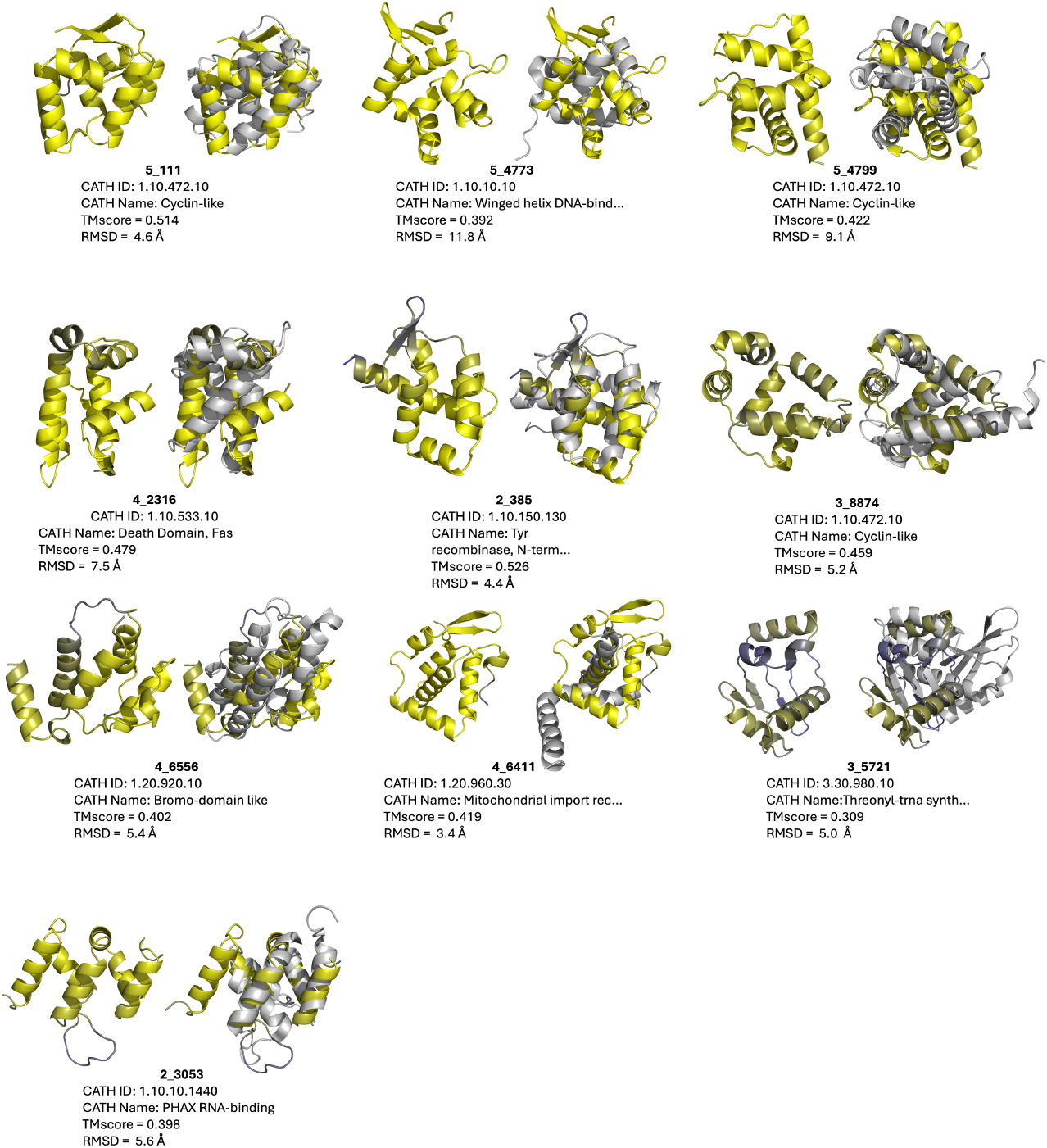
Ten out of 100 novel folds achieved by structure-first foldtuning. Within each pair: **left** putative novel fold (colored by ESMFold pLDDT; yellow=high, blue=low); **right** superimposed with closest CATHDB50 Foldseek hit in TMalign mode, with CATH metadata and global alignment metrics reported below.

**Figure 5.**
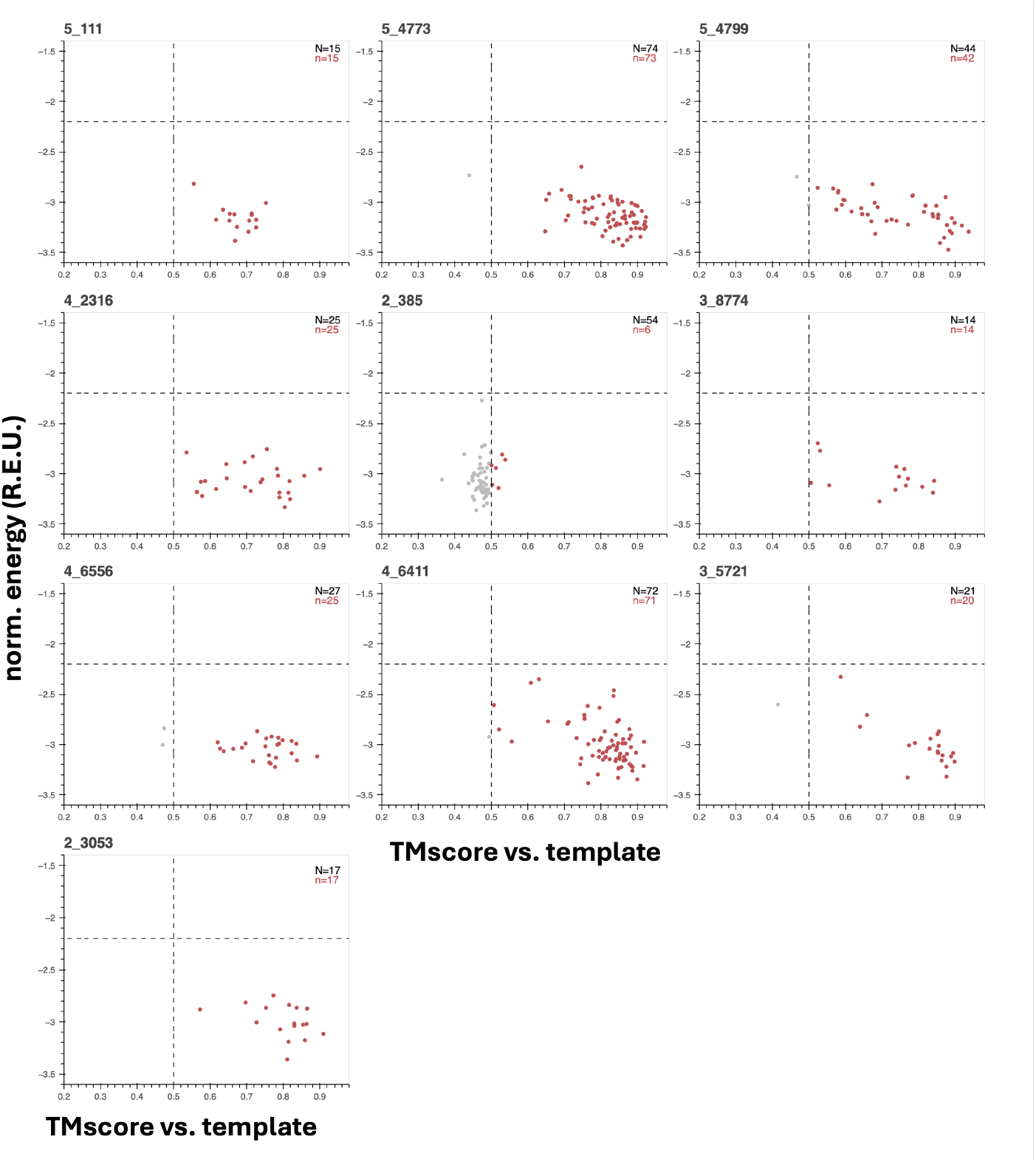
“Inverse-folding landscapes” for ten out of 100 novel folds achieved by structure-first foldtuning imply high stability. Length-normalized energies (from Rosetta) vs. TM-score (from Foldseek in TMalign mode) for ProteinMPNN-designed sequences inverse-folded off of structure-first foldtuning putative novel folds as templates. Gray dots correspond to all sequences/structures for a given template after clustering 200 initial sequences per template at 60% sequence similarity. Red dots show the subset of inverse-folded seqUences whose ESMFold-predicted structures pass an energy scoring threshold (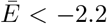 REU/aa) and the standard TM-score global match threshold (TMscore *>* 0.5).

One example, variant 2_385 appears spurious, with a TMscore = 0.526 hit to CATH 1.10.150.130 and an inverse-folding landscape littered with “metastable” analogs with sub-0.5 TMscores upon alignment to the foldtuning-emitted template, suggesting that it is not novel, but a noised version of the natural tyrosine recombinase N-terminal domain (Figs. 4-5, Table S.2). The remaining nine variants, by contrast, impute high stability *in silico*, with strong funnel-esque association between lower-energy folded-states and high TMscore alignments to their putative novel templates, plus nearly all inverse-folded versions clearing the rough energy targets of *< −*2.2 REU/aa and TMscore *>* 0.5 (Fig. 5). By eye, TMscore, and RMSD, these nine are clearly distinct from their closest CATH counterparts and are distributed across all-*α* (5_4799, 4_2316, 3_8774, 4_6556, 2_3053), *α* + *β* (5_4773, 4_6411), and *α/β* (5_111, 3_5721) topologies (Fig. 4). Altogether, this constitutes strong evidence that structure-first foldtuning is able to target novel protein structures with meaningful fitness- and topology-agnostic selection criteria, extracting new-to-nature domains with broad shape diversity from a PLM by steering with synthetic sequences that impart supersecondary structural innovation.

## 3 Discussion

The extent and precise forms of folded domains with structures unlike any of the contents of natural protein-space has been the subject of much debate in structural biology and biophysics. To search for such domains in a concerted fashion, we effectuated two distinct methods for probing new-to-nature regions of protein structure-space using information captured by protein language models (PLMs). In one, we filled out fold-space from scratch using an evolutionary algorithm steered by PLM-driven estimates of sequence and structure reasonableness. In the second, we directly enriched for structural novelty in freely-generated PLM output. The rates of fold discovery – roughly 1-in-200 for the evolutionary algorithm and 1-in-50 for the direct enrichment approach (“structure-first foldtuning”) – are more than 1000x the rate of new fold detection from segmenting and searching *>* 200 million AlphaFoldDB predictions.

All of these efforts used structure prediction models and structure-based search methods; a compelling explanation for the rapid fold emergence rates in our approaches is the application of selection and/or enrichment criteria that reward globularity and compactness without more explicit guidance on secondary, supersecondary, or tertiary structure properties, and without any consideration of functional information whatsoever. What is evident too, is the ability of PLMs as agents of a novelty-first design philosophy, empowered by a capacity to credibly evaluate sequence motifs and – implicitly – structure motifs that emanate from different generative rules than the operative ones of nature. Fascinatingly, prioritizing novelty in structure brings along novelty of sequence as a byproduct – only 1/15 genetic algorithm novel domain candidates and 10/1018 structure-first foltuning novel domain candidates are predicted off of sequences with any statistically significant homolog in UniRef50; inverse-folding off these templates maintains sequence novelty, with only 12/1143 (1.05%) and 21/3912 (0.05%) hits against natural sequence databases, respectively. This suggests that both PLM-based methods have discovered pockets of joint sequence-structure novelty well outside of nature. Our findings align squarely with the position that permissible structure-space is much broader than that covered by nature; and that, conjecturing a step further, there may exist numerous fold ensembles sufficient for the essential processes of life, arising or not subject initial conditions and/or population size effects.

## Acknowledgments and Disclosure of Funding

We thank Steve Mayo, Erik Winfree, Richard Murray, Zach Martinez, Alec Lourenço, Shichen Liu, Joe Boktor, Sonia Yuan, and Hunter Richards, as well as all members of the Thomson Lab for thoughtful comments and discussion.

This work was supported by the National Institutes of Health under award number R01GM150125, the David & Lucille Packard Foundation under a Packard Fellowship to M.T., and the Heritage Medical Research Institute. The authors have no competing interests to disclose.

## 4 Supplemental Material

### 4.1 Methods

#### 4.1.1 Fragment Library Assembly for Genetic Algorithm

The initial fragment library for the structure discovery genetic algorithm was assembled via a replica-exchange Markov Chain Monte Carlo (RE-MCMC) approach for sampling over a sequence landscape. We construct a two-term energy function that combines a preference for accepting mutations that increase sequence-likelihood and a preference for increasing predicted structural contact density, metrics that are inferred from the ESM2-650M model [Lin et al., 2023]. For a *single chain*: if an individual sequence *s*_*i*_ of length *N* has an associated log-likelihood 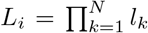 where the *l*_*k*_ represent independent residue-wise likelihoods, we can write Δ*E* for a proposed move from sequence *s*_*i*_ to *s*_*j*_ as

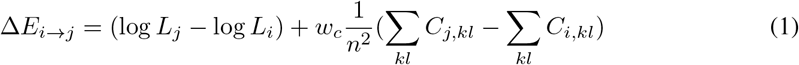

where *n* is the fixed sequence length, *C*_*i*_, *C*_*j*_ are binary contact matrices s.t. *C*_*i,kl*_ = 1 indicates that residues *k* and *l* of sequence *i* are predicted to be in physical contact within *<* 8 Åin the corresponding three-dimensional structure, and *w*_*c*_ controls the relative weights of the likelihood and contact terms. Contact matrices are inferred from the contact_prediction head of ESM2-650M, simultaneously with log-likelihood calculation. We restrict allowable moves to single point mutations, though the formulation holds generally for any *s*_*i*_, *s*_*j*_ of the same length.

The standard Metropolis-Hastings criterion is used, setting the acceptance probability for a proposed single point mutation (sampled uniformly across sequence positions and amino-acid identities) from *s*_*i*_ to *s*_*j*_ for a *single chain* to

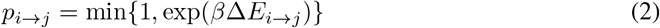

For RE-MCMC, several chains are monitored simultaneously following the above, sampling the same landscape at different temperatures, thereby balancing riskier less-local moves by “hot” chains with more conservative local moves by “cold” chains. Adjacent chains in the temperature array attempt to swap positions on the landscape (and their respective sequences) periodically at a stochastic frequency *λ*; the proposed swap move between chains *i, j* is accepted with probability

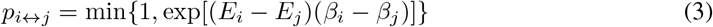

where *{β*_*i*_*}* refers to thermodynamic *β*, the inverse of the sampling temperature *T* .

A total of 800 fragments were generated, running for *n* = 5000 steps, stochastically attempting to swap a uniformly randomly selected pair of adjacent random chains at a rate of *λ* = 0.01 swp/step, 5 chains with inverse temperatures 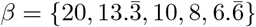 from “cold” to “hot,” and *w*_*c*_ = 1. Initial sequences for all chains {*s*_0_} were random amino-acid strings (sampled uniformly with respect fo amino-acid identity) of length 40; the coldest chain (*β* = 20) sequence at step 5000 was added to the library.

#### 4.1.2 Structure Discovery Genetic Algorithm

The structure discovery genetic algorithm begins by sampling an initial population *P*_0_ of 100 fragments from a fragment library assembled as described. For a fixed number of rounds, the *k*-th round proceeds by:

1. Generating 20 new variants from *P*_*k−*1_. A pair of variants is generated by drawing two sequences uniformly at random from *P*_*k−*1_, performing a crossover operation with the number of crossover points *n*_*cross*_ *∼* Poisson(*λ* = 1.535) and the locations of the crossover points uniformly distributed over the sequence length(s), and performing a mutation operation with the number of mutations *n*_*mut*_ *∼* Binom(*n*_*i*_, *λ* = 0.05) and mutation locations and identities uniformly distributed over sequence lengths.
2. Evolving the new (*k*)-th round variants through eeMHMC with the energy function found in Eq. 1 for *n* = 5000 steps, with *w*_*c*_ = 1, *β* = 10, and adaptive temperature adjustment at 100-step intervals.
3. Adding the evolved variants to *P*_*k−*1_ to form *P*_*k*_.
4. Predicting structures and computing the amino-acid surface-area burial fraction for *all* sequences in *P*_*k*_.
5. Stochastic selection for burial fraction, maintaining a target constant population size of 100.

The above procedure repeats up to a desired number of generations (200 in this study). To enforce constant population size while stochastically eliminating sequences from the population, we note that:

If the number of surviving sequences after round *k* is to be |*P*_*k*_| = *N*_*sel*_, then the expectation of *N*_*sel*_ must be

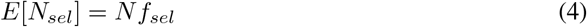

where *N* is the temporary population size after new variants have been added but before any have been removed, and *f*_*sel*_ is the fraction of sequences that are to survive. We can additionally write that

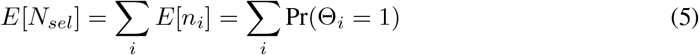

where Θ_*i*_ *∼* Bernoulli(*p*_*i*_) for some mathematically appropriate *p*_*i*_, as the survival of a given sequence is independent of the survival probability of all others. We have the choice of the form of *p*_*i*_ and so take *p*_*i*_ = exp[*−β*_*k*_(0.8*−γ*_*i*_)], where *β*_*k*_ is a sampling hyperparameter to be determined and *γ*_*i*_ is the burial fraction of sequence *s*_*i*_

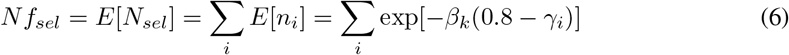

Note that if *γ*_*i*_ *>* 0.8, then this would imply *p*_*i*_ *>* 1; formally, we ought to say *p*_*i*_ = max {1, exp[*−β*_*k*_(0.8*−γ*_*i*_)]}; empirically, however, the burial fraction for even exceptionally well-packed and folded protein domains is bounded above by *γ*_*i*_ = 0.8.

Making the simplifying assumption that the {*γ*_*i*_}’s are roughly normally distributed – empirically justified for ESM2-generated sequence and reasonably extrapolated to sequences being evolved subject to an ESM2-based energy function – we can say

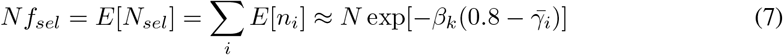

where 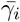 is the mean of all calculated burial fractions in the temporarily augmented population *P*_*k*_ leaving only algebraic rearrangement to solve for our lone sampling hyperparameter *β*_*k*_, effectively a selection inverse temperature, as follows

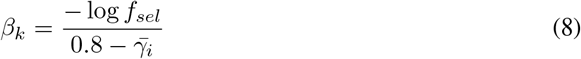

This completes the material necessary to specify and implement the structure discovery genetic algorithm.

#### 4.1.3 Structure-First Foldtuning

Foldtuning was performed and implemented essentially as described in Subramanian et al. [2025] with the following modifications: (1) generation of 10,000 sequences per round in batches of 250; (2) selection of sequences satisfying structural compactness (amino-acid surface burial fraction *>* 0.5) and novelty (no CATHDB50 hit with TMscore *>* 0.5) criteria; (3) ranking of filtered, validated round *n* sequences for round *n* + 1 finetuning in descending order of amino-acid surface burial fraction; (4) omission of the initial “evotuning” round due to absence of a specific target fold.

#### 4.1.4 Selection of Novel Folds for Computational Characterization

For the genetic algorithm experiment, all fifteen putative novel folds were advanced to the computational validation and characterization. For the foldtuning-based experiment, 1018 putative novel folds were initially cumulatively identified over five rounds of structure-first foldtuning. To remove redundancy, predicted structures of the 1018 were clustered with Foldseek at a similarity threshold of TMscore = 0.5, decreasing the number of templates to 916. Given that the structural diversity of the whole AlphaFoldDB50 runs deeper than that of the CATHDB50 subset, the 916 remaining putative novel folds were searched, again using Foldseek, against the entire AlphaFoldDB50, dropping structures with any single hit with alignment region TMscore *>* 0.5. This reduced the number of templates to 762. These 762 templates were ranked in order of decreasing surface-area burial fraction and the top 100 carried through for inverse-folding and energy scoring validation. For Fig. 4 and Fig. 5, only the further top 10 of these top 100, as ranked by lowest (most-stable) mean Rosetta-scored energy over all inverse-folded sequences are depicted.

#### 4.1.5 Structure Prediction and Assignment

All structures were predicted with default ESMFold inference parameters as in Lin et al. [2023]. Predicted structures were annotated to CATH domain labels via Foldseek structure-based search against the prebuilt CATHDB50 database of Lau et al. [2024], running in accelerated TMalign mode. The consensus CATH domain was defined as the fold accounting for the most hits with TMscore *>* 0.5 and max(query_coverage, target_coverage) *>* 0.8. In the absence of at least one hit satisfying these criteria, a structure was considered to be un-assignable.

#### 4.1.6 Basic Chemical Property Calculations

Amino-acid surface area burial fraction was calculated using custom code and reference individual amino-acid surface areas (HMS Bionumbers: 103239).

#### 4.1.7 Energy Scoring Calculations

Biomolecule energy scores were obtained using the default ‘ref2015’ energy function and standard relaxation and scoring workflow in Rosetta v3.11, as described in Alford et al. [2017]. Energy scores are reported in **R**osetta **E**nergy **U**nits (R.E.U.), normalized to sequence length.

#### 4.1.8 Validation of Inverse-Folding Sequences and Structures

For both the genetic algorithm and foldtuning-based experiments, 200 sequences were generated per structural template with ProteinMPNN, using the vanilla—v_48_020 model, sampling temperature 0.2, Gaussian backbone noise with per-coordinate *σ* = 0.1 Å, and forced omission of the rare/ambiguous amino acids B, J, O, U, X, and Z [Dauparas et al., 2022]. Within each batch of 200, sequences were downclustered at 60% sequence identity with mmseqs2, structures predicted with ESMFold, and queried against the template structure with Foldseek in TMalign mode using the standard TMscore *>* 0.5 threshold as confirmation of a global match.

### 4.2 Supplemental Figures

**Figure S1:**
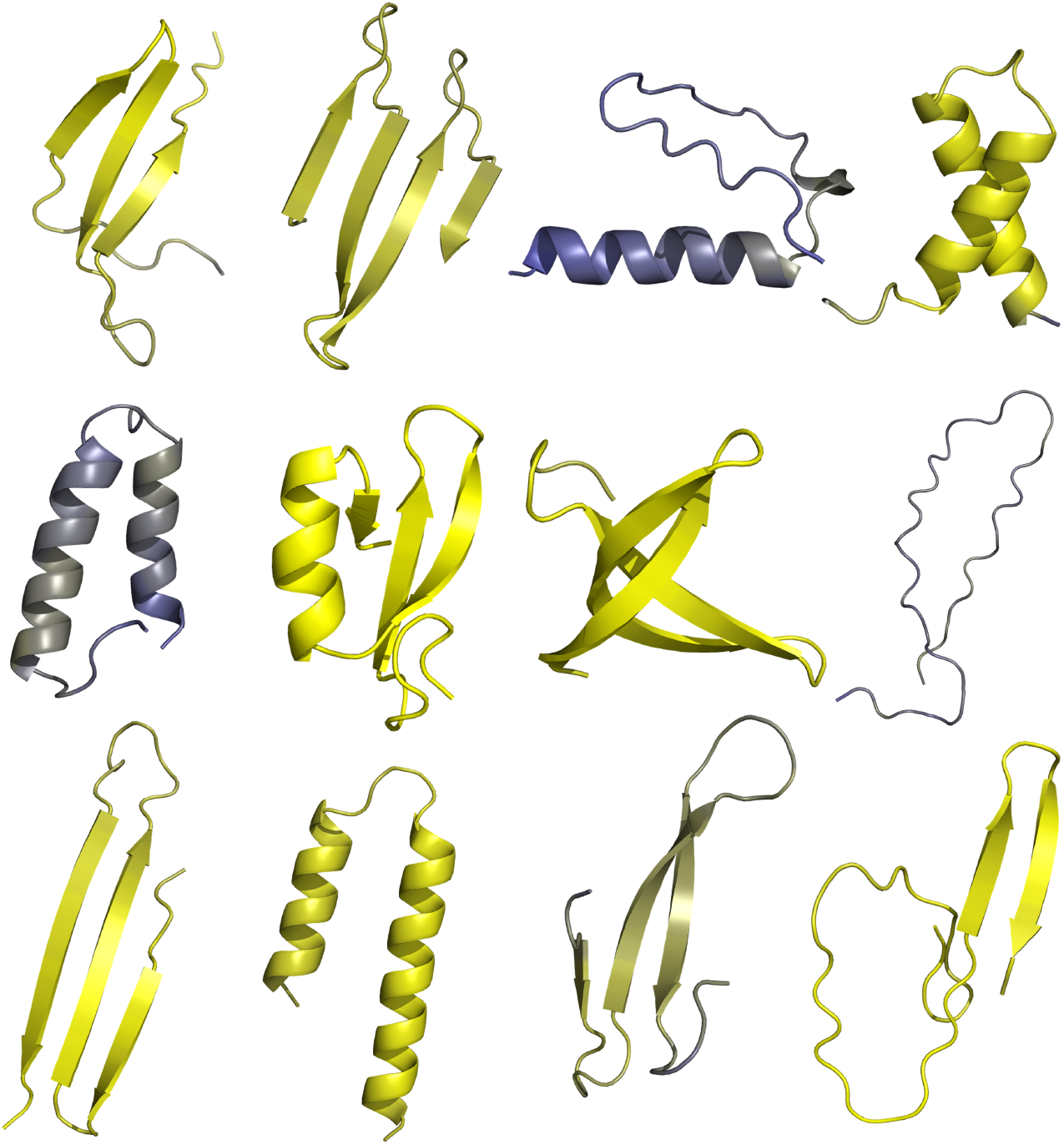
Example structure fragments generated by RE-eeMHMC. 10 of 800 structure fragments predicted from sequences designed by replica-exchange explore-exploit Metropolis-Hastings Monte Carlo sampling (RE-eeMHMC). Individual structures are colored by ESMFold pLDDT; yellow=high, blue=low.

### 4.3 Supplemental Tables

**Table S.1:**
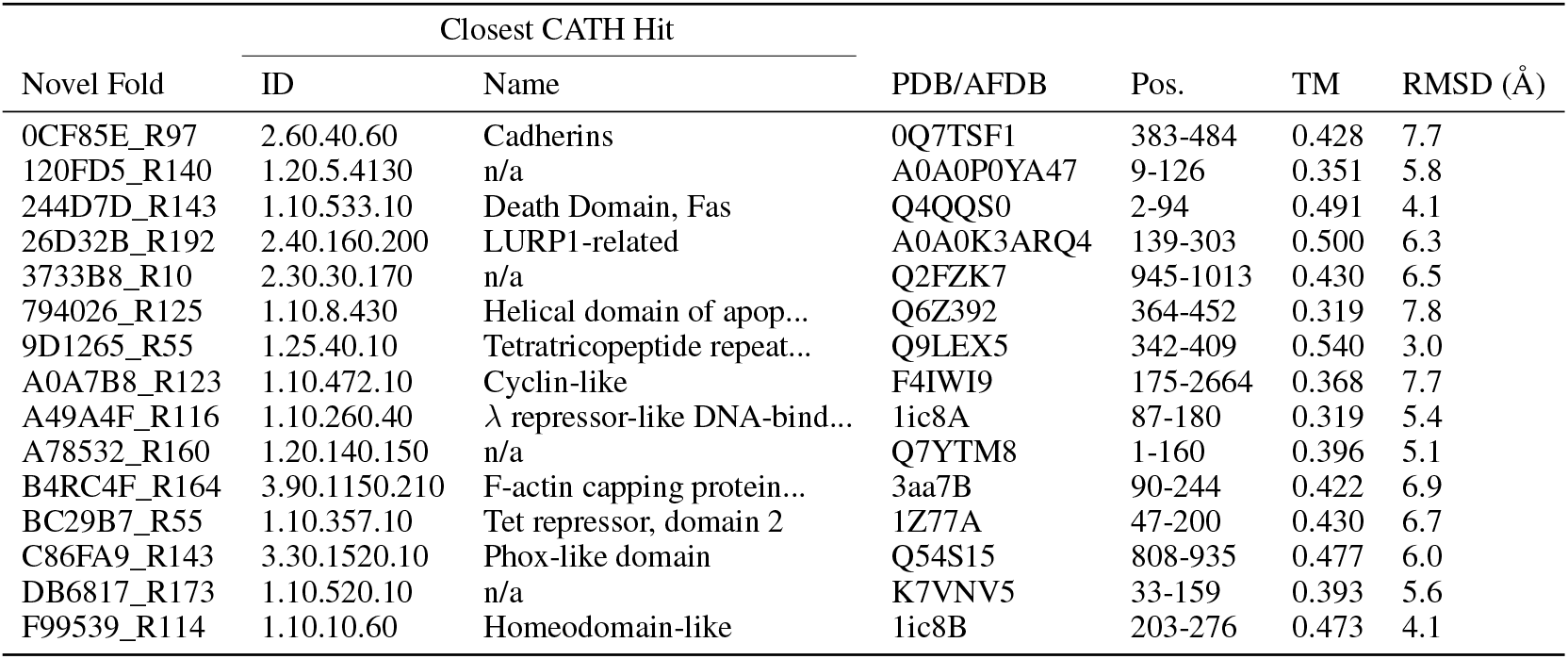
Metadata and structural alignment metrics for closest CATH domain Foldsheek hits to 15 novel folds proposed by the genetic algorithm approach.

**Table S.2:**
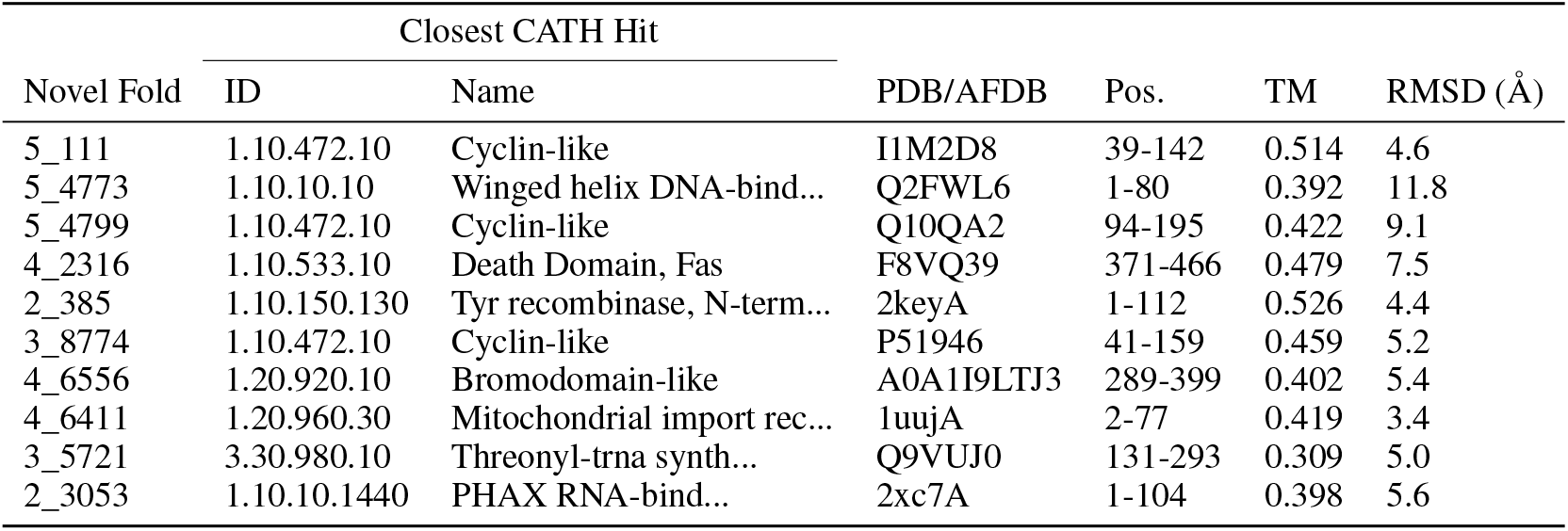
Metadata and structural alignment metrics for closest CATH domain Foldsheek hits to 10 novel folds proposed by structure-first foldtuning.

## References

R. F. Alford, A. Leaver-Fay, J. R. Jeliazkov, M. J. O’Meara, F. P. DiMaio, H. Park, M. V. Shapovalov, P. D. Renfrew, V. K. Mulligan, K. Kappel, J. W. Labonte, M. S. Pacella, R. Bonneau, P. Bradley, R. L. J. Dunbrack, R. Das, D. Baker, B. Kuhlman, T. Kortemme, and J. J. Gray. The Rosetta AllAtom Energy Function for Macromolecular Modeling and Design. Journal of Chemical Theory and Computation, 13(6):3031–3048, June 2017.ISSN 1549-9618. doi: 10.1021/acs.jctc.7b00125. URL https://doi.org/10.1021/acs.jctc.7b00125. Publisher: American Chemical Society.

C. Alvarez-Carreño, R. J. Gupta, A. S. Petrov, and L. D. Williams. Creative destruction: New protein folds from old. Proceedings of the National Academy of Sciences, 119(52):e2207897119. Dec. 2022. doi: 10.1073/pnas.2207897119. URL https://www.pnas.org/doi/abs/10.1073/pnas.2207897119. Publisher: Proceedings of the National Academy of Sciences.

I. Anishchenko, S. J. Pellock, T. M. Chidyausiku, T. A. Ramelot, S. Ovchinnikov, J. Hao, K. Bafna, C. Norn, A. Kang, A. K. Bera, F. DiMaio, L. Carter, C. M. Chow, G. T. Montelione, and D. Baker. De novo protein design by deep network hallucination. Nature, 600 (7889):547–552, Dec. 2021.ISSN 1476-4687. doi: 10.1038/s41586-021-04184-w. URL https://www.nature.com/articles/s41586-021-04184-w. Number: 7889 Publisher: Nature Publishing Group.

D. Baker. A surprising simplicity to protein folding. Nature, 405(6782):39–42, May 2000.ISSN 1476-4687. doi: 10.1038/35011000. URL https://www.nature.com/articles/35011000. Number: 6782 Publisher: Nature Publishing Group.

I. Barrio-Hernandez, J. Yeo, J. Jänes, M. Mirdita, C. L. M. Gilchrist, T. Wein, M. Varadi, S. Velankar, P. Beltrao, and M. Steinegger. Clustering predicted structures at the scale of the known protein universe. Nature, 622(7983):637–645, Oct. 2023.ISSN 1476-4687. doi: 10.1038/s41586-023-06510-w. URL https://www.nature.com/articles/s41586-023-06510-w. Number: 7983 Publisher: Nature Publishing Group.

E. Bornberg-Bauer. How are model protein structures distributed in sequence space? Biophysical Journal, 73(5):2393–2403, Nov. 1997. ISSN 00063495. doi: 10.1016/S0006-3495(97)78268-7. URL https://linkinghub.elsevier.com/retrieve/pii/S0006349597782687.

B. Chen, X. Cheng, Y.-a. Geng, S. Li, X. Zeng, B. Wang, J. Gong, C. Liu, A. Zeng, Y. Dong, J. Tang, and L. Song. xTrimoPGLM: Unified 100B-Scale Pre-trained Transformer for Deciphering the Language of Protein. Technical report, bioRxiv, July 2023. URL https://www.biorxiv.org/content/10.1101/2023.07.05.547496v3. Section: New Results Type: article.

B. Chitturi, S. Shi, L. N. Kinch, and N. V. Grishin. Compact Structure Patterns in Proteins. Journal of Molecular Biology, 428(21):4392–4412, Oct. 2016.ISSN 0022-2836. doi: 10.1016/j.jmb.2016.07.022. URL https://www.sciencedirect.com/science/article/pii/S0022283616302893.

I.-G. Choi and S.-H. Kim. Evolution of protein structural classes and protein sequence families. Proceedings of the National Academy of Sciences, 103(38):14056–14061, Sept. 2006. doi: 10.1073/pnas.0606239103. URL https://www.pnas.org/doi/abs/10.1073/pnas.0606239103. Publisher: Proceedings of the National Academy of Sciences.

J. Dauparas, I. Anishchenko, N. Bennett, H. Bai, R. J. Ragotte, L. F. Milles, B. I. M. Wicky, Courbet, R. J. de Haas, N. Bethel, P. J. Y. Leung, T. F. Huddy, S. Pellock, D. Tischer, F. Chan, Koepnick H. Nguyen, A. Kang, B. Sankaran, A. K. Bera, N. P. King, and D. Baker. Robust deep learning–based protein sequence design using ProteinMPNN. Science, 378(6615):49–56, Oct. 2022. doi: 10.1126/science.add2187. URL https://www.science.org/doi/10.1126/science.add2187. Publisher: American Association for the Advancement of Science.

K. A. Dill and H. S. Chan. From Levinthal to pathways to funnels. Nature Structural Biology, 4 (1):10–19, Jan. 1997.ISSN 1545-9985. doi: 10.1038/nsb0197-10. URL https://www.nature.com/articles/nsb0197-10. Publisher: Nature Publishing Group.

C. L. Dupont, A. Butcher, R. E. Valas, P. E. Bourne, and G. Caetano-Anollés. History of biological metal utilization inferred through phylogenomic analysis of protein structures. Proceedings of the National Academy of Sciences, 107(23):10567–10572, June 2010. doi: 10.1073/pnas.0912491107. URL https://www.pnas.org/doi/abs/10.1073/pnas.0912491107. Publisher: Proceedings of the National Academy of Sciences.

J. Durairaj, A. M. Waterhouse, T. Mets, T. Brodiazhenko, M. Abdullah, G. Studer, G. Tauriello, M. Akdel, A. Andreeva, A. Bateman, T. Tenson, V. Hauryliuk, T. Schwede, and J. Pereira. Uncovering new families and folds in the natural protein universe. Nature, 622(7983):646–653, Oct. 2023.ISSN 1476-4687. doi: 10.1038/s41586-023-06622-3. URL https://www.nature.com/articles/s41586-023-06622-3. Publisher: Nature Publishing Group.

N. Ferruz, S. Schmidt, and B. Höcker. ProtGPT2 is a deep unsupervised language model for protein design. Nature Communications, 13(1):4348, July 2022.ISSN 2041-1723. doi: 10.1038/s41467-022-32007-7. URL https://www.nature.com/articles/s41467-022-32007-7. Number: 1 Publisher: Nature Publishing Group.

S. Govindarajan and R. A. Goldstein. Why are some proteins structures so common? Proceedings of the National Academy of Sciences, 93(8):3341–3345, Apr. 1996. doi: 10.1073/pnas.93.8.3341. URL https://www.pnas.org/doi/abs/10.1073/pnas.93.8.3341. Publisher: Proceedings of the National Academy of Sciences.

T. Hayes, R. Rao, H. Akin, N. J. Sofroniew, D. Oktay, Z. Lin, R. Verkuil, V. Q. Tran, J. Deaton, M. Wiggert, R. Badkundri, I. Shafkat, J. Gong, A. Derry, R. S. Molina, N. Thomas, Y. A. Khan, C. Mishra, C. Kim, L. J. Bartie, M. Nemeth, P. D. Hsu, T. Sercu, S. Candido, and A. Rives. Simulating 500 million years of evolution with a language model, Dec. 2024. URL https://www.biorxiv.org/content/10.1101/2024.07.01.600583v2. Pages: 2024.07.01.600583 Section: New Results.

R. Helling, H. Li, R. Mélin, J. Miller, N. Wingreen, C. Zeng, and C. Tang. The designability of protein structures. Journal of Molecular Graphics and Modelling, 19(1):157–167, Feb. 2001.ISSN 1093-3263. doi: 10.1016/S1093-3263(00)00137-6. URL https://www.sciencedirect.com/science/article/pii/S1093326300001376.

B. L. Hie, K. K. Yang, and P. S. Kim. Evolutionary velocity with protein language models predicts evolutionary dynamics of diverse proteins. Cell Systems, 13(4):274–285.e6, Apr. 2022.ISSN 2405-4712, 2405-4720. doi: 10.1016/j.cels.2022.01.003. URL https://www.cell.com/cell-systems/abstract/S2405-4712(22)00038-2. Publisher: Elsevier.

R. S. Kim, E. Levy Karin, M. Mirdita, R. Chikhi, and M. Steinegger. BFVD—a large repository of predicted viral protein structures. Nucleic Acids Research, 53(D1):D340–D347, Jan. 2025a.ISSN 1362-4962. doi: 10.1093/nar/gkae1119. URL https://doi.org/10.1093/nar/gkae1119.

W. Kim, M. Mirdita, E. Levy Karin, C. L. M. Gilchrist, H. Schweke, J. Söding, E. D. Levy, and M. Steinegger. Rapid and sensitive protein complex alignment with Foldseek-Multimer. Nature Methods, 22(3):469–472, Mar. 2025b.ISSN 1548-7105. doi: 10.1038/s41592-025-02593-7. URL https://www.nature.com/articles/s41592-025-02593-7. Publisher: Nature Publishing Group.

B. Kuhlman, G. Dantas, G. C. Ireton, G. Varani, B. L. Stoddard, and D. Baker. Design of a Novel Globular Protein Fold with Atomic-Level Accuracy. Science, 302(5649):1364–1368, Nov. 2003. doi: 10.1126/science.1089427. URL https://www.science.org/doi/full/10.1126/science.1089427. Publisher: American Association for the Advancement of Science.

A. M. Lau, N. Bordin, S. M. Kandathil, I. Sillitoe, V. P. Waman, J. Wells, C. A. Orengo, and D. T. Jones. Exploring structural diversity across the protein universe with The Encyclopedia of Domains. Science, 386(6721):eadq4946. Nov. 2024. doi: 10.1126/science.adq4946. URL https://www.science.org/doi/10.1126/science.adq4946. Publisher: American Association for the Advancement of Science.

H. Li, R. Helling, C. Tang, and N. Wingreen. Emergence of Preferred Structures in a Simple Model of Protein Folding. Science, 273(5275):666–669, Aug. 1996. doi: 10.1126/science.273.5275.666. URL https://www.science.org/doi/abs/10.1126/science.273.5275.666. Publisher: American Association for the Advancement of Science.

Z. Lin, H. Akin, R. Rao, B. Hie, Z. Zhu, W. Lu, N. Smetanin, R. Verkuil, O. Kabeli, Y. Shmueli, A. dos Santos Costa, M. Fazel-Zarandi, T. Sercu, S. Candido, and A. Rives. Evolutionary-scale prediction of atomic-level protein structure with a language model. Science, 379(6637):1123–1130, Mar. 2023. doi: 10.1126/science.ade2574. URL https://www.science.org/doi/10.1126/science.ade2574. Publisher: American Association for the Advancement of Science.

L. M. Longo, D. Despotović, O. Weil-Ktorza, M. J. Walker, J. Jabłonśka, Y. Fridmann-Sirkis, G. Varani, N. Metanis, and D. S. Tawfik. Primordial emergence of a nucleic acid-binding protein via phase separation and statistical ornithine-to-arginine conversion. Proceedings of the National Academy of Sciences, 117(27):15731–15739, July 2020. doi: 10.1073/pnas.2001989117. URL https://www.pnas.org/doi/abs/10.1073/pnas.2001989117. Publisher: Proceedings of the National Academy of Sciences.

A. Madani, B. Krause, E. R. Greene, S. Subramanian, B. P. Mohr, J. M. Holton, J. L. Olmos, C. Xiong, Z. Z. Sun, R. Socher, J. S. Fraser, and N. Naik. Large language models generate functional protein sequences across diverse families. Nature Biotechnology, pages 1–8, Jan. 2023.ISSN 1546-1696. doi: 10.1038/s41587-022-01618-2. URL https://www.nature.com/articles/s41587-022-01618-2. Publisher: Nature Publishing Group.

S. Minami, N. Kobayashi, T. Sugiki, T. Nagashima, T. Fujiwara, R. Tatsumi-Koga, G. Chikenji, and N. Koga. Exploration of novel-protein folds through de novo design. Nature Structural & Molecular Biology, pages 1–9, July 2023.ISSN 1545-9985. doi: 10.1038/s41594-023-01029-0. URL https://www.nature.com/articles/s41594-023-01029-0. Publisher: Nature Publishing Group.

G. Munsamy, S. Lindner, P. Lorenz, and N. Ferruz. ZymCTRL: a conditional language model for the controllable generation of artificial enzymes. NeurIPS Machine Learning for Structural Biology Workshop, Dec. 2022. URL https://www.mlsb.io/papers_2022/ZymCTRL_a_conditional_language_model_for_the_controllable_generation_of_artificial_enzymes.pdf.

G. A. Pavlopoulos, F. A. Baltoumas, S. Liu, O. Selvitopi, A. P. Camargo, S. Nayfach, A. Azad, S. Roux, L. Call, N. N. Ivanova, I. M. Chen, D. Paez-Espino, E. Karatzas, I. Iliopoulos, K. Konstantinidis, J. M. Tiedje, J. Pett-Ridge, D. Baker, A. Visel, C. A. Ouzounis, S. Ovchinnikov, A. Buluç, and N. C. Kyrpides. Unraveling the functional dark matter through global metagenomics. Nature, 622(7983):594–602, Oct. 2023.ISSN 1476-4687. doi: 10.1038/s41586-023-06583-7. URL https://www.nature.com/articles/s41586-023-06583-7. Number: 7983 Publisher: Nature Publishing Group.

K. Sakuma, N. Kobayashi, T. Sugiki, T. Nagashima, T. Fujiwara, K. Suzuki, N. Kobayashi, T. Murata, T. Kosugi, R. Tatsumi-Koga, and N. Koga. Design of complicated all- protein structures. Nature Structural & Molecular Biology, pages 1–8, Jan. 2024.ISSN 1545-9985. doi: 10.1038/s41594-023-01147-9. URL https://www.nature.com/articles/s41594-023-01147-9. Publisher: Nature Publishing Group.

J. Skolnick, H. Zhou, and M. Brylinski. Further Evidence for the Likely Completeness of the Library of Solved Single Domain Protein Structures. The Journal of Physical Chemistry B, 116(23): 6654–6664, June 2012.ISSN 1520-6106. doi: 10.1021/jp211052j. URL https://doi.org/10.1021/jp211052j. Publisher: American Chemical Society.

A. M. Subramanian, Z. A. Martinez, A. L. Lourenço, S. Liu, and M. Thomson. Unexplored regions of the protein sequence-structure map revealed at scale by a library of foldtuned language models, Jan. 2025. URL https://www.biorxiv.org/content/10.1101/2023.12.22.573145v2. Pages: 2023.12.22.573145 Section: New Results.

W. R. Taylor, V. Chelliah, S. M. Hollup, J. T. MacDonald, and I. Jonassen. Probing the “Dark Matter” of Protein Fold Space. Structure, 17(9):1244–1252, Sept. 2009.ISSN 0969-2126. doi: 10.1016/j.str.2009.07.012. URL https://www.cell.com/structure/abstract/S0969-2126(09)00295-0. Publisher: Elsevier.

M. Varadi, S. Anyango, M. Deshpande, S. Nair, C. Natassia, G. Yordanova, D. Yuan, O. Stroe, G. Wood, A. Laydon, A. Žídek, T. Green, K. Tunyasuvunakool, S. Petersen, J. Jumper, E. Clancy, R. Green, A. Vora, M. Lutfi, M. Figurnov, A. Cowie, N. Hobbs, P. Kohli, G. Kleywegt, E. Birney, D. Hassabis, and S. Velankar. AlphaFold Protein Structure Database: massively expanding the structural coverage of protein-sequence space with high-accuracy models. Nucleic Acids Research, 50(D1):D439–D444, Jan. 2022.ISSN 0305-1048. doi: 10.1093/nar/gkab1061. URL https://doi.org/10.1093/nar/gkab1061.

R. Verkuil, O. Kabeli, Y. Du, B. I. M. Wicky, L. F. Milles, J. Dauparas, D. Baker, S. Ovchinnikov, T. Sercu, and A. Rives. Language models generalize beyond natural proteins. Technical report, bioRxiv, Dec. 2022. URL https://www.biorxiv.org/content/10.1101/2022.12.21.521521v1. Section: New Results Type: article.

P. Vyas, O. Trofimyuk, L. M. Longo, F. K. Deshmukh, M. Sharon, and D. S. Tawfik. Helicase-like functions in phosphate loop containing beta-alpha polypeptides. Proceedings of the National Academy of Sciences, 118(16):e2016131118. Apr. 2021. doi: 10.1073/pnas.2016131118. URL https://www.pnas.org/doi/abs/10.1073/pnas.2016131118. Publisher: Proceedings of the National Academy of Sciences.

J. L. Watson, D. Juergens, N. R. Bennett, B. L. Trippe, J. Yim, H. E. Eisenach, W. Ahern, A. J. Borst, R. J. Ragotte, L. F. Milles, B. I. M. Wicky, N. Hanikel, S. J. Pellock, A. Courbet, W. Sheffler, J. Wang, P. Venkatesh, I. Sappington, S. V. Torres, A. Lauko, V. De Bortoli, E. Mathieu, S. Ovchinnikov, R. Barzilay, T. S. Jaakkola, F. DiMaio, M. Baek, and D. Baker. De novo design of protein structure and function with RFdiffusion. Nature, pages 1–3, July 2023.ISSN 1476-4687. doi: 10.1038/s41586-023-06415-8. URL https://www.nature.com/articles/s41586-023-06415-8. Publisher: Nature Publishing Group.

A. L. Watters, P. Deka, C. Corrent, D. Callender, G. Varani, T. Sosnick, and D. Baker. The Highly Cooperative Folding of Small Naturally Occurring Proteins Is Likely the Result of Natural Selection. Cell, 128(3):613–624, Feb. 2007.ISSN 0092-8674, 1097-4172. doi: 10.1016/j.cell.2006.12.042. URL https://www.cell.com/cell/abstract/S0092-8674(07)00117-1. Publisher: Elsevier.

K. Yue and K. A. Dill. Forces of tertiary structural organization in globular proteins. Proceedings of the National Academy of Sciences, 92(1):146–150, Jan. 1995. doi: 10.1073/pnas.92.1.146. URL https://www.pnas.org/doi/abs/10.1073/pnas.92.1.146. Publisher: Proceedings of the National Academy of Sciences.

Y. Zhang, I. A. Hubner, A. K. Arakaki, E. Shakhnovich, and J. Skolnick. On the origin and highly likely completeness of single-domain protein structures. Proceedings of the National Academy of Sciences, 103(8):2605–2610, Feb. 2006. doi: 10.1073/pnas.0509379103. URL https://www.pnas.org/doi/abs/10.1073/pnas.0509379103. Publisher: Proceedings of the National Academy of Sciences.

